# M-Risk: A framework for assessing global fisheries management efficacy of sharks, rays, and chimaeras

**DOI:** 10.1101/2022.05.29.493344

**Authors:** C. Samantha Sherman, Glenn Sant, Colin A. Simpfendorfer, Eric D. Digel, Patrick Zubick, Grant Johnson, Michael Usher, Nicholas K. Dulvy

**Affiliations:** Earth to Oceans Research Group, Department of Biological Sciences, Simon Fraser University, Burnaby, British Columbia, Canada; TRAFFIC International, Cambridge, U.K.; Australian National Centre for Ocean Resources and Security, University of Wollongong, NSW, Australia; Institute of Marine and Antarctic Studies, University of Tasmania, Hobart, Tasmania, Australia; Department of Industry, Tourism and Trade, Fisheries Branch, Northern Territory Government, Darwin, Northern Territory, Australia

**Keywords:** Data-poor fisheries, ecological risk assessment, elasmobranch, marine conservation, resource management, socio-ecological resilience

## Abstract

Fisheries management is essential to guarantee sustainable capture of target species and avoid undesirable declines of incidentally captured species. A key challenge is halting and reversing declines of shark and ray species, and specifically assessing the degree to which management is sufficient to avoid declines in relatively data-poor fisheries. While ecological risk analyses focus on intrinsic ‘productivity’ and extrinsic ‘susceptibility’, one would ideally consider the influence of ‘fisheries management’. Currently, there is no single management evaluation that can be applied to a combination of fishery types at the scale of individual country or Regional Fisheries Management Organisations (RFMOs). Here, we outline a management risk (M-Risk) framework for sharks, rays, and chimaeras used to evaluate species’ risk to overfishing resulting from ineffective management. We illustrate our approach with application to one country (Ecuador) and RFMO (Inter-American Tropical Tuna Commission) and illustrate the variation in scores among species. We found that while both management units assessed had similar overall scores, the scores for individual attributes varied. Ecuador scored higher in reporting-related attributes, while the IATTC scored higher in attributes related to data collection and use. We evaluated whether management of individual species was sufficient for their relative sensitivity by combining the management risk score for each species with their intrinsic sensitivity to determine a final M-Risk score. This framework can be applied to determine which species face the greatest risk of overfishing and be used by fisheries managers to identify effective management policies by replicating regulations from countries with lower risk scores.

## 1. INTRODUCTION

Catch in marine fisheries globally has decreased since the mid-1990s despite increasing effort (FAO, 2020; Pauly, Zeller, & Palomares, 2021; Rousseau, Watson, Blanchard, & Fulton, 2019). Ensuring long-term sustainable fisheries requires management not only of target species but also of all other species affected by the fishery (Hilborn et al., 2003). However, halting biodiversity loss and ecosystem management ideally requires knowledge of the status and fishing mortality associated with all species taken in the fishery. Globally, at least 13,060 different species are caught (FAO, 2021), leaving over 97% of species without a stock assessment – there are only 957 stock assessments for 360 unique species (RAM Legacy Stock Assessment Database, 2021). For such data-rich stock assessed populations and species, we know their levels of exploitation and their sustainable fishing mortality levels (RAM Legacy Stock Assessment Database, 2021; Ricard, Minto, Jensen, & Baum, 2012). But for data-poor species, how do we determine their status? Much of our understanding on data-poor species’ catch is based on landings data reported to the Food and Agriculture Organization of the United Nations (FAO)(Froese, Zeller, Kleisner, & Pauly, 2012). Data-poor species comprise >80% of global catch and almost two-thirds of species are overexploited (Costello et al., 2012; Guan, Chen, Boenish, Jin, & Shan, 2020). However, landings data are not consistent or reliable globally, particularly in countries with many landing sites and/or high levels of artisanal catch, particularly for sharks and their relatives (Khan et al., 2020; Okes & Sant, 2022; Ruano-Chamorro, Subida, & Fernández, 2017). Additionally, the catches of data-poor species, if characterised, are often reported as aggregates; grouped together by genus, family, or higher, giving little information on the catch of individual species (Cashion, Bailly, & Pauly, 2019; FAO, 2019). The portfolio effects of these aggregate catch statistics mask serial depletions, cryptic declines, and local extinctions (Dulvy, Metcalfe, Glanville, Pawson, & Reynolds, 2000; Lawson et al., 2020; Schindler et al., 2010). Many methods have been developed to assess data-poor species and stocks, which require, *inter alia*, different data inputs such as catches (Anderson, Branch, Ricard, & Lotze, 2012), length compositions (Cope & Punt, 2009; Froese, 2004), age-at-maturity (Brooks, Powers, & Cortés, 2010; Cope, 2013), gear selectivity (Le Quesne & Jennings, 2012). However, all have major assumptions and biases associated with the outputs (Chrysafi & Kuparinen, 2015). There remains a need for a rapid risk assessment technique, when detailed stock assessments are not possible, at the scale of countries and RFMOs.

When assessing the risk of a species within a fishery, the terminology used throughout the risk assessment literature is inconsistent. Therefore, we have used the most common terms and defined them where appropriate. Here, we use the framing of Vulnerability as derived from the social science, hazard assessment, and climate risk literature (Turner II et al., 2003), and increasingly used in biological risk assessment (Allison et al., 2009; Williams, Shoo, Isaac, Hoffmann, & Langham, 2008). Vulnerability is typically considered to be the interaction of intrinsic sensitivity (biological traits that inform extinction risk) with exposure (the overlap between a threat and the species’ range) to a threatening process (i.e., fishing or climate change). Broadly, vulnerability can be considered to be the inverse of ecological resilience (Allison et al., 2009). Taken together, the Potential Impact (sensitivity x exposure) can be offset either by phenotypical plasticity or genotypic evolution when considering a species, or through building Adaptive Capacity of the human system, which can be strengthened through management or disaster planning and preparedness (Allen Consulting Group, 2005; Allison et al., 2009; Dulvy et al., 2011). For example, fishers in Kenya increased their Adaptive Capacity to climate change by increasing community infrastructure and access to credit (Cinner et al., 2015). Ecological risk assessments can take several different forms, however, all measure similar aspects of risk such that a species’ vulnerability (*V*) is a function of their intrinsic life history sensitivity (Intrinsic Sensitivity), Exposure to fishing pressure, say as inferred from the spatial and depth overlap with the species’ distribution, and the species’ catchability in the fishing gears (Hobday et al., 2011; Walker et al., 2021). All this Potential Impact can be offset by Adaptive Capacity:

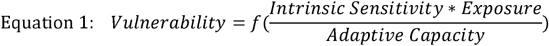

This generic risk assessment framework has been pruned down to focus only on the Potential Impact, specifically the interplay of sensitivity and exposure. This is most commonly treated by Productivity Susceptibility Assessments (PSA), a type of ecological risk analysis which has been typically applied to single fishery to compare the relative risk of the full range incidentally captured species (Fletcher, 2005; Hobday et al., 2011; Micheli, De Leo, Butner, Martone, & Shester, 2014) (**Figure 1**). This approach can be applied to data-poor fisheries and in highly diverse systems with taxonomically diverse species in the incidental catch, such as in an Australian Prawn Trawl Fishery (Astles et al., 2006; Stobutzki, Miller, & Brewer, 2001).

**Figure 1.**
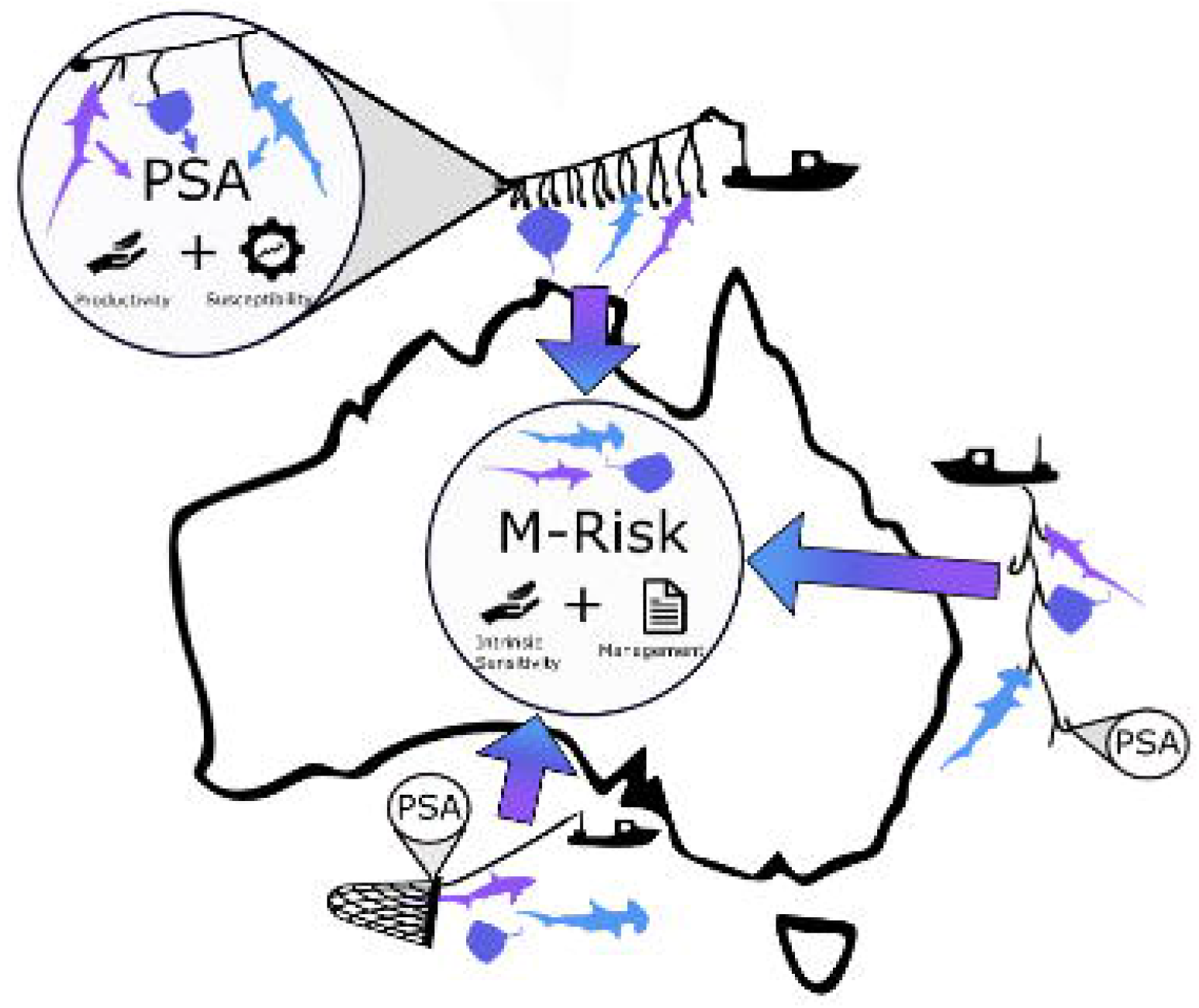
The goal of M-Risk assessments is to determine relative species risk within and across fisheries at country and RFMO scales. Whereas Productivity Susceptibility Analysis (PSA) is applied to a specific assessment unit (i.e., typically the fishery), and hence it is not typical to make comparisons among PSAs. PSAs identify Vulnerability based on a species’ attributes (Productivity) and its overlap with the fishery being assessed (Susceptibility). M-Risk also identifies Vulnerability based on a species’ attributes (albeit with a different name: Intrinsic Sensitivity), however, this M-Risk is contingent on its management within the fishery, not just spatial overlap with the fishery.

As with all risk assessments, there are trade-offs to achieve a result that best informs the goals of the assessment. Other risk frameworks, like PSAs, tend to focus on a specific assessment unit comprised of a single fishery, set of species, or gear type, or combination (Astles et al., 2006; Cortés, Brooks, & Shertzer, 2015; Zhou, Milton, & Fry, 2012). PSA customization allows for a deeper, more nuanced assessment within a single assessment unit. However, specificity of a PSA makes comparisons difficult to other PSAs with different criteria or scoring. Thus, the need for a method that can be used across all fishery types including different gear types, target species, and data availability at the scale of countries and RFMOs. Similarly, while a risk assessment can be used to rank the species at risk within a fishery, it does not consider the degree to which the Potential Impact is or can be offset by the existence of some form of management (Turner II et al., 2003) (**Figure 1**).

To incorporate management into risk assessments, a rapid management risk assessment (M-Risk) was proposed for shark species and tested for species with high intrinsic sensitivity, many of which are CITES listed (Lack, Sant, Burgener, & Okes, 2014). The results showed variation in management efficacy despite protections that should have resulted from compliance with CITES listings. While the original risk framework was fit for the purpose of threatened species, it was not necessarily optimised for species caught and traded in high volumes. M-Risk considers that vulnerability (*V*) is a function of a species’ intrinsic sensitivity (*IS*), exposure to fishing pressure (*E*), and Management, substituted for Adaptive Capacity from equation 1:

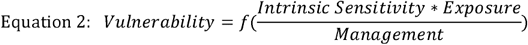

We note that while management does not influence intrinsic susceptibility *per se* it does influence the interaction of intrinsic sensitivity and exposure *inter alia* by managing the availability, encounterability, selectivity, and post-release mortality of species through various technical approaches (Hobday et al., 2011).

Sharks, rays, and chimaeras (class: Chondrichthyes; >1,199 species, hereafter “sharks and rays”) present a challenge for fisheries managers as they are frequently retained as secondary catch or discarded as incidental catch (Stevens, Bonfil, Dulvy, & Walker, 2000). Overfishing, both targeted and incidental, has caused populations of sharks and rays to decline dramatically over the past 50 years, leading to a high rate of elevated extinction risk (Dulvy et al., 2021; Pacoureau et al., 2021). As they are commonly considered unavoidable incidental or secondary catch, sharks and rays are frequently treated as unimportant or unavoidable in a management context. Therefore, sharks and rays are an ideal candidate group for an M-Risk assessment. Many sharks and rays are longer-lived species, with low intrinsic rates of population increase, and cannot be fished at the same rate as most teleosts, despite often being caught alongside them (Brander, 1981; Musick, Burgess, Cailliet, Camhi, & Fordham, 2000; Myers & Worm, 2005; Pardo, Cooper, Reynolds, & Dulvy, 2018; Stevens, Walker, & Simpfendorfer, 1997). Fisheries management of sharks and rays is further complicated because of the diversity of species, gears used, jurisdictions in which they are caught (Dulvy et al., 2017; Simpfendorfer & Dulvy, 2017), and their complex migration patterns that transit the waters of multiple countries, which increases their exposure in multiple fisheries and cumulative impacts of being caught throughout their migration routes (Dragičević, Dulčić, & Capapé, 2009; Heupel et al., 2015; Sellas et al., 2015). Despite the high intrinsic sensitivity of many species, sustainable shark and ray fisheries are possible if assessment and management is adequate. There are 39 sustainably fished populations of 33 species of sharks and rays around the globe (Simpfendorfer & Dulvy, 2017) and signs of recovery in US and EU managed populations (Amelot et al., 2021; Peterson et al., 2017). Sustainable shark and ray fisheries all have common characteristics that distinguish them from unsustainable shark and ray fisheries. These centre on fishing mortality to sustainable levels through limits on catch and/ or effort, supported by robust legislation, well-enforced regulations, and science-based advisory processes (Simpfendorfer & Dulvy, 2017; Woodhams, Peddemors, Braccini, Victorian Fisheries Authority, & Lyle, 2021).

Here, we provide a revised set of attributes based on the original M-Risk framework (Lack et al. 2014), for application to any shark or ray species in any fishery to rapidly and objectively assess management efficacy in a comparable consistent manner at country and RFMO scales (**Figure 1**). First, we illustrate the revised M-Risk framework with two case studies: one for all species in a country (Ecuador, 29 species) and one for 24 species in a Regional Fisheries Management Organisation (RFMO) – the Inter-American Tropical Tuna Commission (IATTC). Second, once assessments were completed, management scores for each species were combined with their intrinsic risk scores to attain a final M-Risk score per species in each management unit. Third, we discuss the methodology and initial assessments with the goal of completing assessments for a larger suite of management units globally, hence species mentioned includes some not assessed in this paper. These scores are intended for two anticipated uses: (1) by fisheries managers so that improvements can be made specifically for species that are undermanaged relative to their intrinsic sensitivity or for overall fisheries improvements (risk management) and (2) for comparative analysis of the state of the world’s shark and ray fisheries.

## 2. DEVELOPING THE NEW M-RISK FRAMEWORK

### 2.1 Unit of Assessment – country and RFMO

When considering exploited marine species and their management, there are two levels at which an assessment can be completed: stock or fishery. First, a stock is usually a political construct that consists of a demographically isolated portion of the global population that is often managed as a single unit (Begg & Waldman, 1999). Where a single stock is widespread, the fisheries may be managed separately by different jurisdictions. This can complicate the risk assessment of the stock, as it may not be clear which management unit is responsible for the species’ status if it is not uniformly co-managed across jurisdictions, and the assessment must factor in the effects of multiple sectors and the cumulative impacts on the status of the stock (Urquhart, Acott, Symes, & Zhao, 2014). For example, the Winter Skate (*Leucoraja ocellata*) occurs in both Canadian and USA waters; however, the population trend in the USA is increasing, while the population trend in Canadian waters is decreasing (Kulka et al., 2020). The discrepancy may be due to differing management focus and effectiveness of the fisheries in each country (Kulka et al., 2020). Second, in our case, a fishery is defined by its management such that each separate fishery is managed by a single jurisdiction that regulates the laws of operation within the spatial extent of the fishery.

We are applying M-Risk at two scales (country and RFMO), hereafter referred to as “Management Units”. Within a single country (Ecuador, here), we selected a representative fishery based on which one had the highest level of catch (retained and discarded) with the species under assessment. Each fishery will be assessed as a single unit, not considering if the species is caught in other fisheries within or outside of the management unit, or the cumulative impacts of multiple fisheries contributing to fishing mortality of the species. High seas fishing was not considered when assessing countries, only fishing occurring within their respective Exclusive Economic Zones (EEZs), as high seas management should be picked up in RFMO assessments. Vessels from member countries of each RFMO are obligated to adhere to the RFMO regulations when operating in those fishery grounds. Countries may apply stricter regulations to their flagged vessels fishing within RFMO grounds. However, we are applying our criteria only to legislation that is applicable for **all** member countries of the RFMO, as this is the minimum standard in those fishing grounds.

### 2.2 Species Selection

Sharks and rays are commonly caught and traded globally (Dent & Clarke, 2015). As there are currently over 1,200 described species (Ebert, Dando, & Fowler, 2021; Last et al., 2016), we curated a list consisting of the most frequently traded species worldwide, based on four sources: (i) catches reported to the Food and Agriculture Organization of the United Nations (FAO) (FAO, 2019), (ii) reconstructed catches from the Sea Around Us Project (Pauly & Zeller, 2016), (iii) species listed on the Appendices of the Convention on International Trade in Endangered Species of Wild Fauna and Flora (CITES) (https://cites.org), and (iv) species listed on the Appendices of the Convention on the Conservation of Migratory Species of Wild Animals (CMS) (https://www.cms.int). Additionally, we included species or groups that are caught in high abundances but usually identified at a higher taxonomic level, such as cowtail rays (*Pastinachus* spp.) in the Indo-Pacific. The final list for the full M-Risk project included 69 sharks identified to species level apart from one genus (*Etmopterus;* n = 38), 27 rays identified to either species level or a ray group (e.g., eagle rays) (16 species; 11 groups, n = 80), and 6 ghost shark species for a total of 102 species or groups (**Data S1**). The ray groups included different levels of taxonomic resolution for species that were difficult to distinguish, have similar life history characteristics, distribution, and human use, or are taxonomically unresolved or may comprise a species complex (e.g., maskrays – *Neotrygon* spp.). The number of species assessed in this paper was lower, at 36 species (30 sharks, 2 ray species, and 4 ray groups). However, the larger suite of 102 species is referred to here because intrinsic risk scores of the 36 species considered here are expressed relative to the entire 102 species of the project. Assessments were completed for each species separately, except for the groups, for which a single assessment was completed for the whole group. The combined assessments were necessary, as there are very limited species-specific data or management system details within the groups. All species that have a distribution overlapping with the management unit’s spatial grounds are assessed for that management unit. Final M-Risk scores were calculated at the species level, as intrinsic sensitivity of species within each group differed.

### 2.3 Management Assessment Framework

We devised 21 measurable attributes to assess the degree to which management is adequate to prevent sharks and rays from being overexploited. These attributes were chosen based on regulations that (1) enabled understanding, and (2) curbed fishing mortality on the focal shark and ray species and was informed by previous work on sustainable shark fisheries (Davidson, Krawchuk, & Dulvy, 2015; FAO, 1999; Melnychuk, Peterson, Elliott, & Hilborn, 2017). Some attributes were considered but could not be included due to data paucity. For example, total catch of each species is not available in most fisheries and, therefore, was not included as an attribute at present but could be considered as more of these data become available.

Twenty-one attributes were created comprised of three common classes and two classes related to the spatial unit of analysis (**Table 1**):

**Table 1.**
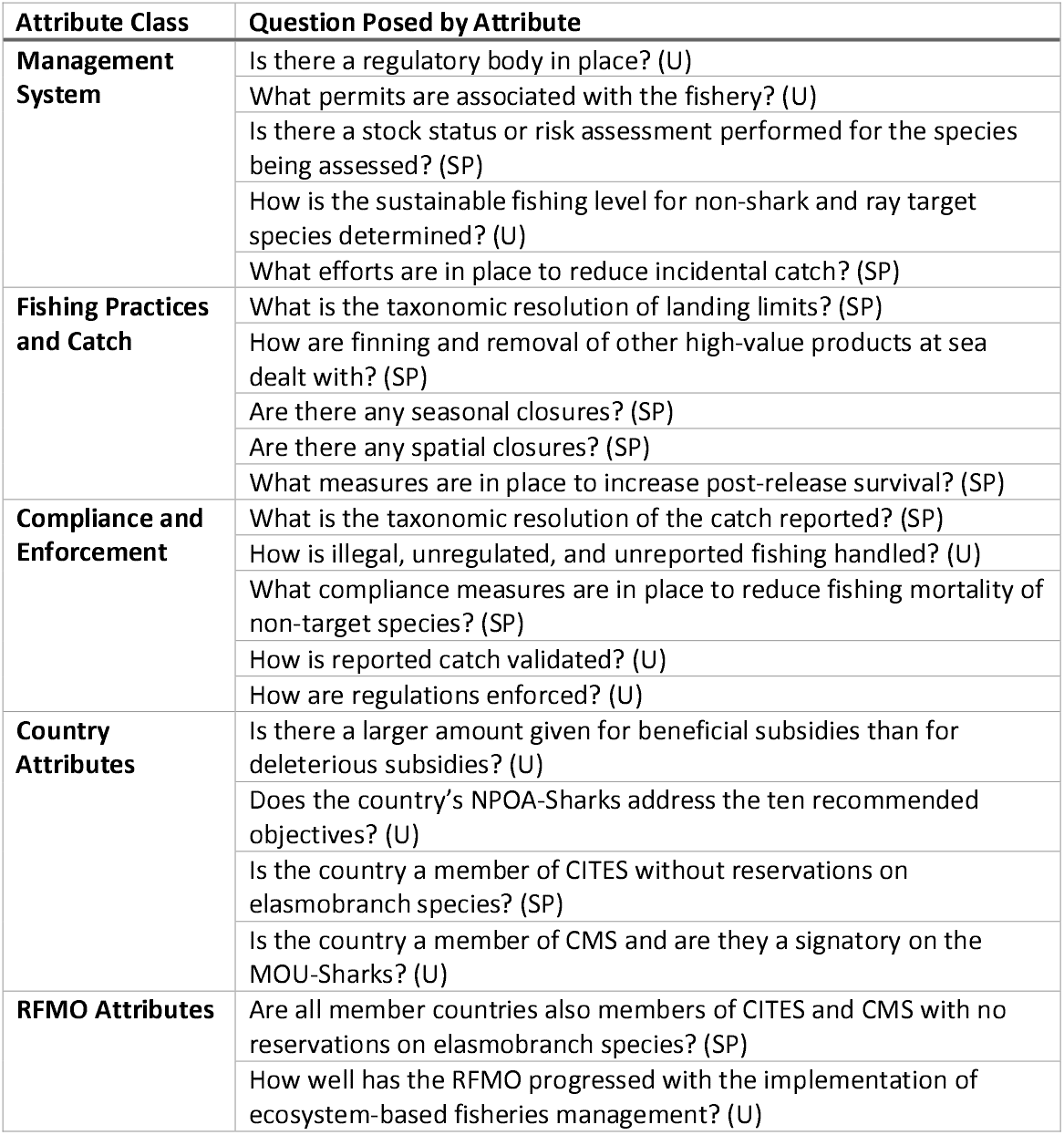
The 21 M-Risk Attributes used for assessments. Universal attributes are indicated by a (U) and species-specific attributes are indicated by a (SP). For complete explanation and scoring of each attribute see **supplementary information 1**.

i. he management system (5 attributes),
ii. fishing practices and catch (5 attributes),
iii. compliance and enforcement (5 attributes), and an additional category considered the management unit and how it relates to other management units which consisted of attributes either specific to:
iv. country (4 attributes), OR
v. RFMO (2 attributes).

Therefore, countries were scored for 19 of the attributes and RFMOs are scored for 17 of the attributes. These numbers were different as there were more aspects of a country that could be assessed. Next, we unpack each of the five attribute classes.

#### 2.3.1 Management System

The management system consists of a regulatory body, fishery permits, and assessments to understand the potential risk the fishery poses. Without such foundational management in place, it is unlikely the fishery will have any more sophisticated regulations or any capacity to enforce regulations in place. These attributes are intended to increase our understanding of the capability of management for improvement. Having basic management, such as a permitting system, enables fisheries managers to have records of all vessels operating in the fishery, which can enable the setting of limits on catches or effort. For management units that receive low scores in the ‘management system’ attributes, we would recommend the government allocates resources to strengthen the capacity of the fishery to manage the resource as a first step to introducing intricate regulations that are more likely to curb fishing mortality and increase sustainability.

#### 2.3.2 Fishing Practices and Catch

Fishing practices and catch include landing limits, shark finning, post-release survival, and closures. Management of at-sea fishing operations is difficult due to the size of the area in which fishing occurs and the costs associated with patrolling large spaces (Rowlands, Brown, Soule, Boluda, & Rogers, 2019). Ideally, fisheries managers would know exactly the number of each species caught and their fate post-release (if released), in addition to having spatial and/or seasonal closures in place for at-risk species at sensitive times or locations. Simply, having regulations in place that affect at-sea operations but can be measured or enforced upon landing are the most effective, as these do not require boarding of vessels at-sea. For example, the requirement that sharks are landed with fins naturally attached is something that can be checked upon landing the catch (Fowler & Séret, 2010). Similarly, the existence of spatial and temporal closures can be monitored through use of Vessel Monitoring Systems (VMS) that can be checked from land (Enguehard, Devillers, & Hoeber, 2013). For management units with low scores in the ‘fishing practices and catch’ attributes, we would recommend the inclusion of regulations that decrease fishing mortality on intrinsically sensitive species, like sharks and rays.

#### 2.3.3 Compliance, Monitoring, and Enforcement

Compliance, monitoring, and enforcement consists of reporting catch, ensuring reports are valid, having enforcement in place for violations, and monitoring of illegal, unregulated, and unreported fishing (IUU). Compliance, meaning operating within the established legislation, often requires fisher agreement. Fishers are more likely to comply with legislation (1) when they understand regulations are in place for their own interest by increasing sustainability, (2) through feeling obligated to comply, and (3) through enforcement measures that may decrease their profitability if non-compliance is discovered (Hønneland, 1999). Additionally, fishers with a better relationship and trust with the management authority are more likely to comply with regulations (Hauck, 2008). In an ideal world, fishers would have 100% compliance with the regulations, however, this is not realistic, and therefore, must be monitored and enforced (Price et al., 2016). The attributes within this class were designed to assess how the quantity and composition of the catch is validated and what measures are in place to ensure there is high compliance within the fishery. Similarly, we include an attribute considering how IUU fishing is accounted for through management measures. As IUU fishing has been estimated to be up to 26 million tonnes per year of the global catch, this can have a significant impact on fishing mortality and must be accounted for in assessing and managing the fishery (Agnew et al., 2009). The attributes within this group require more nuanced regulations, therefore, management units with higher scores in this category are expected to have higher scores in the previous categories as well. For management units with lower scores in the ‘compliance, monitoring, and enforcement’ attributes, we would recommend allocating funding to increase fishery officer coverage, including at landing ports to observe unloading and in office to monitor vessel operations.

#### 2.3.4 Country Attributes

Country attributes include whether the country is involved in international agreements, how effective their National Plan of Action for Sharks (NPOA-Sharks) is, if one exists, and the amount given in subsidies. These attributes deal with the relationship of the country to the rest of the world through the engagement with international treaties and agreements like CITES and CMS. The scores in this group are indicative of how managers in each country consider preservation of at-risk species. For countries with lower scores in the country attributes, we would recommend improving or drafting an NPOA-Sharks, and reallocating subsidy funding to more beneficial outcomes like marine protected area implementation and management as opposed to tax exemptions and fuel subsidies.

#### 2.3.5 RFMO Attributes

RFMO attributes include whether membership parties are involved in international treaties and agreements like CITES and CMS, and their progress on ecosystem-based fisheries management (EBFM). Similar to the country attribute, membership in CITES and CMS shows willingness to protect species-at-risk and EBFM progress shows the care of the RFMO for the overall ecosystem in which they are fishing. For RFMOs with lower scores in these attributes, we would recommend improving their progress towards EBFM and encouraging member countries to participate in CITES and CMS, without species reservations.

### 2.4 Scoring of Attributes

Each attribute was scored individually based on detailed value statements (e.g., **Table 2**), such that a higher score indicated a higher likelihood of achieving a sustainable outcome for the species (full value statements available in **Supplementary Information 1**). Our scoring of attributes assigned the highest scores to the ideal ‘counterfactual’ situation (Juan-Jordá, Murua, Arrizabalaga, Dulvy, & Restrepo, 2018). We chose narrow ranges for ordinal scoring, (either 0-3, 0-4, or 0-5, similar to a Likert scale) and scored attributes against the value statements found in **supplementary information 1**. In cases where information was absent, precautionary scores of zero were given (Hobday et al., 2011). In some cases, an attribute was not applicable to the species being assessed (i.e., an attribute determining post-release survival is not relevant to a species that is always retained). When this occurred, the attribute would simply not be scored and left out of the final calculations. The narrow range of scores ensured consistency of scoring over time, across jurisdictions, and across assessors. As such the range of uncertainty in scoring can only be low. We did consider scoring over a wider scale, say 0-9, which would have allowed assigning a range instead of a single score, and, therefore, allow incorporation and propagation of uncertainty, say with a Fuzzy Logic or Bayesian approach. However, we deemed that it would be much more difficult to ensure consistent scoring across assessors and jurisdictions. For countries, nine of the attributes were universal (U) for the entire fishery and the remaining ten attributes were specific to each species being assessed (SP). For RFMOs, seven were universal and ten were species-specific (**Table 1**). The final management score, when all attributes were assessed, was calculated based on points scored out of potential points available and converted to a percentage towards ideal management.

**Table 2.**
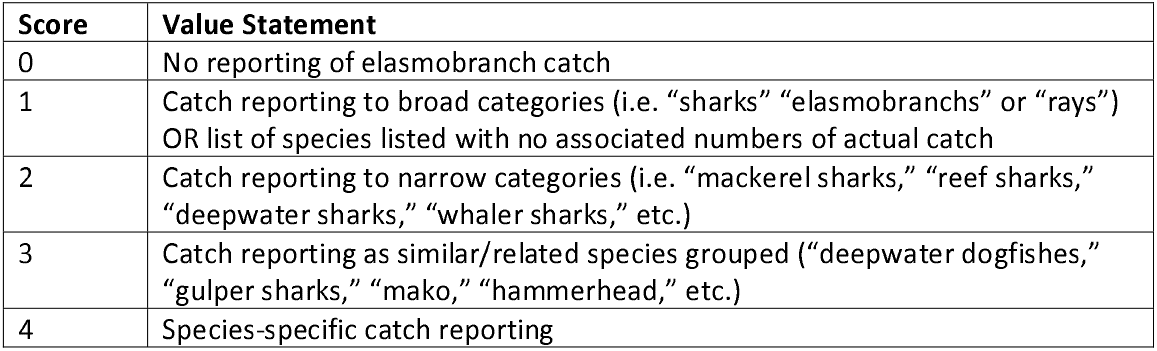
Example of value statements for the attribute that asks, “What is the taxonomic resolution of the catch reported?”

| Score | Value Statement |
| --- | --- |
| 0 | No reporting of elasmobranch catch |
| 1 | Catch reporting to broad categories (i.e. “sharks” “elasmobranchs” or “rays”) OR list of species listed with no associated numbers of actual catch |
| 2 | Catch reporting to narrow categories (i.e. “mackerel sharks,” “reef sharks,” “deepwater sharks,” “whaler sharks,” etc.) |
| 3 | Catch reporting as similar/related species grouped (“deepwater dogfishes,” “gulper sharks,” “mako,” “hammerhead,” etc.) |
| 4 | Species-specific catch reporting |

For RFMOs, scores were assigned based on legislation agreed to by all member countries. Some countries may have additional requirements, however, these were not considered for the RFMO score because we took a precautionary approach and assessed based on the minimum standards for all vessels operating within a fishery. Here, we use the term ‘score’ rather than the term ‘index’. While the academic literature uses the term ‘index’ to refer to any mathematical formula by which different variables are combined, the policy world tends to use the word ‘index’ only when a metric or score has been adopted for use. The reality is that there are many candidate indicators, but very few are fit-for-purpose following extensive performance testing {Newson, 2009 #8482; Rice, 2005 #8483}. The highest possible score (100%) indicates ideal management for sharks and rays; therefore, final management scores for each management unit indicated their progress towards the ideal. Management scores are intended to be used for comparisons between management units and species, and are of little value alone as no fisheries are expected to achieve a score of 100%. Comparisons can be made (1) between different species within a single management unit, (2) the average score between different management units, (3) between a single species in each management unit that it has been assessed, and (4) between different taxonomic groups within or across different management units.

### 2.5 Representing Intrinsic Sensitivity

To understand Vulnerability (equation 2) we calculate the intrinsic sensitivity of a species to overexploitation in addition to the management risk assessment. Factors that contribute to a greater likelihood of population decline or higher intrinsic sensitivity for marine species include large body size and a slow speed of life (e.g. slow somatic growth rate, late age-at-maturity, or long generation length) (Juan-Jordá, Mosqueira, Freire, & Dulvy, 2015; Lee & Jetz, 2011; Reynolds, Dulvy, Goodwin, & Hutchings, 2005).

Elasmobranchs as a group are characterised as having long life spans, late age-at-maturity, and low fecundity (Cortés, 2000; Field, Meekan, Buckworth, & Bradshaw, 2009), and consequently they have a range of intrinsic sensitivities (Pardo, Kindsvater, Reynolds, & Dulvy, 2016). However, within the group there are extremes for all characteristics. For example, the Whale Shark (*Rhincodon typus*) is the largest fish species in the world, attaining a maximum body length of up to 20 m (Chen, Liu, & Joung, 1997). On the other end of the scale, the Dwarf Lanternshark (*Etmopterus perryi*) attains a maximum body length of just 21 cm, approximately one hundredth the maximum body length of Whale Shark (Ebert et al., 2021). Relating to the speed of life, Rigby and Simpfendorfer (2015) discuss the high intrinsic sensitivity of deepwater sharks due to their late age-at-maturity despite their relatively small body size, and thus, the consequences related to their capture in developing deepwater fisheries.

Intrinsic sensitivity can be categorized as low, medium, or high, based on a number of traits that can include age-at-maturity, length-at-maturity, longevity, maximum body length, fecundity, reproductive strategy, and trophic level (Cheung, Pitcher, & Pauly, 2005; Georgeson et al., 2020; Musick, 1999). Data are not available for most shark and ray species for all these traits, therefore we can only use a few reliably available across all species. Oldfield et al. (2012) suggested that minimum age-at-maturity, reproductive strategy, and maximum body length were the three most important factors for sharks and rays, respectively. However, there have been considerable advances in comparative life history theory and it is clear that there are three dimensions to consider in this order of importance: maximum body length, speed of life traits (growth rate, age at maturity, maximum age), and reproductive output (Cortés, 2000; Juan-Jordá, Mosqueira, Freire, & Dulvy, 2013). Many of these traits are not commonly known for shark and ray species and surprisingly, reproductive output contributes little to the maximum intrinsic rate of population increase, except in sharks and rays with very low fecundity of typically fewer than five pups per year (Forrest & Walters, 2009; Pardo et al., 2016), therefore, fecundity is least useful in determining intrinsic sensitivity. Similarly, reproductive strategy would be a binary option, either egg-laying (oviparity) or giving birth to live young (viviparity), and without corresponding information on fecundity or the relationship to maximum intrinsic rate of population increase, would be difficult to directly compare. Finally, trophic level is widely available based on dietary analyses and was considered, however, due to limited species-specific data and the change in trophic level with ontogenetic dietary shifts, we did not include this as a variable when calculating intrinsic sensitivity (Bethea et al., 2007; Lucifora, García, Menni, Escalante, & Hozbor, 2009).

For our analyses, we used two traits to represent intrinsic sensitivity – generation length (the midpoint between age-at-maturity and maximum age) and maximum body length. Many of the shark and ray generation lengths reported in IUCN Red List Assessments are inferred or suspected from closely related species (65% of species considered by M-Risk; **Supplementary Information 2**). Thus, we categorised generation lengths into 5-year bins, such that species with longer generation lengths have higher intrinsic sensitivities. We scored our confidence in the generation length for each species based on whether it was species-specific and the range was within a 5-year bin (high confidence), range spanned multiple 5-year bins or was based on estimated life-history parameters (medium), or was inferred from a congener (low)(**Data S1**). There was a significant positive linear relationship between generation length and relative maximum body length (**Supplementary Information 2**). However, when considering only species with high confidence generation lengths, there was no significant relationship (**Supplementary Information 2 - Figure 1**). This is likely a result of estimated generation lengths being scaled to maximum body length. We then included maximum body size as a second measure of intrinsic sensitivity, of which we could be confident in the species-specific values (**Figure 2**). Ideally, we would use maximum weight but there are few such measures for all species and we encourage their collection. Therefore, we used maximum linear dimension, derived from either maximum length or maximum disc width for some rays, and scored it such that the largest species (*Maximum size*_*L*_) of those we assessed was assigned the reference highest intrinsic sensitivity (i.e., *Relative Size* = 100%) and the *Maximum size*_*x*_ for each other species (*x*) was scaled as a percentage of the largest species (**Figure 2**):

**Figure 2.**
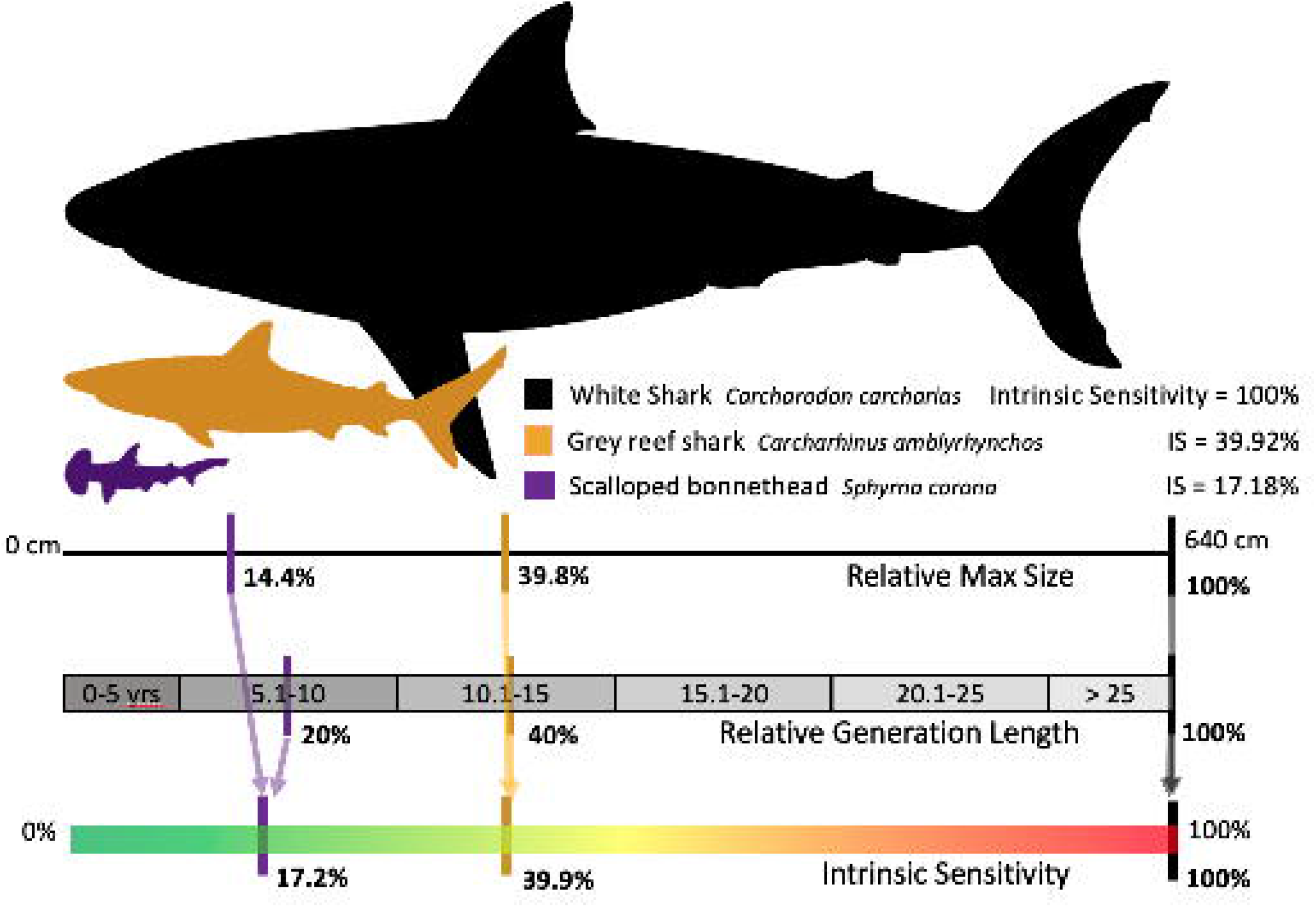
Intrinsic sensitivity (IS) scores based on generation length and maximum size. Higher intrinsic sensitivities are assigned to sharks and rays with longer generation lengths and larger relative sizes. Relative generation length is scored by bins (0-5 years = 0%, 5.1-10 = 20%, 10.1-15 = 40%, 15.1-20 = 60%, 20.1-25 = 80%, >25 = 100%).

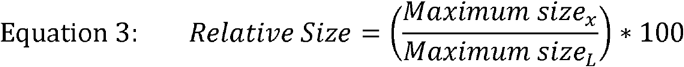

We considered four groups of body morphologies because they are measured in different ways (sharks and shark-like rays, pelagic rays, classic rays, and ghost sharks), and took the largest species for each body morphology and assigned them at 100% *Relative Size*. Whale Shark and Basking Shark (*Cetorhinus maximus*) are outliers, measuring over 1,200 cm longer than the next largest species, therefore, we considered the White Shark (*Carcharodon carcharias*) for these calculations, which attains a maximum size of 640 cm. All these three shark species were assigned a *Relative Size* score of 100% for their maximum size. Intrinsic sensitivity values in this paper are based on the larger project species list (**Data S1**), rather than just those species within this paper to ensure that *Relative Size* is conserved. Both intrinsic sensitivity scores (*Relative GL* and *Relative Size*) were averaged for an overall intrinsic sensitivity (*IS*) score (**Figure 2**):

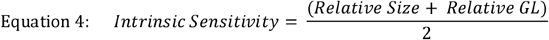

It is not possible for a species to receive an intrinsic sensitivity score of 0% because all species are intrinsically at risk, even if the risk is small.

The intrinsic sensitivity (IS) was then divided by the management assessment score to attain a final M-Risk vulnerability score for each species within each management unit they are assessed:

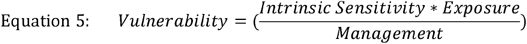

As species exposure to each fishery is difficult to measure, we assume that Exposure = 1 as assessments are only performed on species that are caught or directly affected by the fishery in question. This is the maximum possible exposure and is consistent with the precautionary principle. For countries, there are 19 attribute scores (*A*) with a maximum possible score for each attribute of *Max.A*, and weighting (*W*), therefore, the formula to calculate the M-Risk score is:

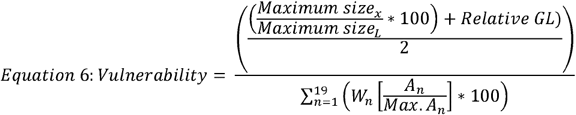

We have weighted all attributes equally (*W* = 1) so the bottom of equation 6 simplifies to the total points scored over the total possible points. The final M-Risk formula simplifies to:

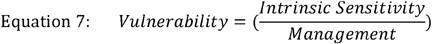

## 3. APPLYING THE M-RISK FRAMEWORK

To explore the application of the M-Risk, we applied our criteria to two case studies, one country (Ecuador), and one RFMO (Inter-American Tropical Tuna Commission, IATTC).

### 3.1 Case Studies: Country and RFMO for Management Assessments

We completed assessments for all species that occur within two management units in order to determine the efficacy of our management-risk assessment framework; one country (Ecuador) and one RFMO (Inter-American Tropical Tuna Commission, IATTC). These assessments were completed for 35 species in Ecuador (23 sharks, 12 rays) and 27 species in the IATTC (20 sharks, 7 rays).

#### 3.1.1 Case Study 1: Ecuador

##### 3.1.1.1 Information Sources

We searched Google and Google Scholar in both English and Spanish for the following four elements in sequence: (i) Ecuadorian fisheries regulations, (ii) specific attribute keywords (i.e., “Ecuador shark finning regulations”), (iii) species name (English, Ecuadorian, Spanish, and Latin binomial) and “Ecuador fishery”, and (iv) species name in all relevant languages and specific attribute keywords (i.e., “Ecuador Whale Shark catch limits”) as has been done previously in ecological risk assessments (Cortés et al., 2010). Ten of the attributes were fishery-specific and did not require steps iii and iv. All attributes were scored on the most current publicly available information. We acknowledge, however, that in some cases this information may be out-of-date, and the most recent regulations are not available online. In cases where no information was available, the precautionary approach was applied, and it was concluded that there were no regulations related to the species in question and a score of zero was given. Again, we acknowledge that this information may exist, but we were unable to access due to it not being publicly available or readily accessible. For this reason, we do not consider uncertainty surrounding scores as exhaustive searches were completed and our framework was developed with the intent to reward transparency, i.e., if we couldn’t find the information in a reasonable amount of time then this was regarded as problematic reflecting lower levels of transparency and availability and thus scored as a zero. The point is that knowledge that exists somewhere, but is not available, is not likely helpful to management or transparency.

Four primary resources were used to complete the Ecuador management risk assessments: (1) Ecuador’s National Fisheries Institute website (www.institutopesca.gob.ec), (2) Ministry of Aquaculture and Fisheries website (acuaculturaypesca.gob.ec), (3) the Ecuador Law website (www.derechoecuador.com), and (4) a paper by Martínez-Ortiz et al. (2015) describing enforcement measures and species-specific catch data. Although multiple fisheries exist in Ecuador, two were chosen to be representative of their management. The Large Pelagic Artisanal Fishery was selected as it accounted for ∼93% of shark and ray catch in Ecuador (Alava, Lindop, & Jacquet, 2015). The only exception was for the Whale Shark as there is common interaction between this species and purse-seine vessels that does not exist with other species, thus the Purse Seine Fishery was chosen to assess Whale Shark management in Ecuador (Dagorn, Holland, Restrepo, & Moreno, 2013; Rowat & Brooks, 2012).

##### 3.1.1.2 Management Assessment Results

Management assessment scores from all 35 species ranged from 53 to 68% of the ideal score. One species (Whale Shark) received the highest score of 68%, five species (3 sharks, 2 rays) scored 67%, and the final 29 species (19 sharks, 10 rays) scored 53 to 62% (**Figure 3a; Table S1**). The differences in management scores were mainly due to three attributes: (1) landing limits, (2) post-release survival, and (3) catch reporting. Scores for the landing limits attribute ranged from 0 to 3 (the highest score possible), while post-release survival had range of 0 to 2 (out of a possible 3) and catch reporting ranged from 1 to 4 (out of 4). The scores for the remaining 16 attributes were the same across all species in the Large Pelagic Artisanal Fishery.

**Figure 3.**
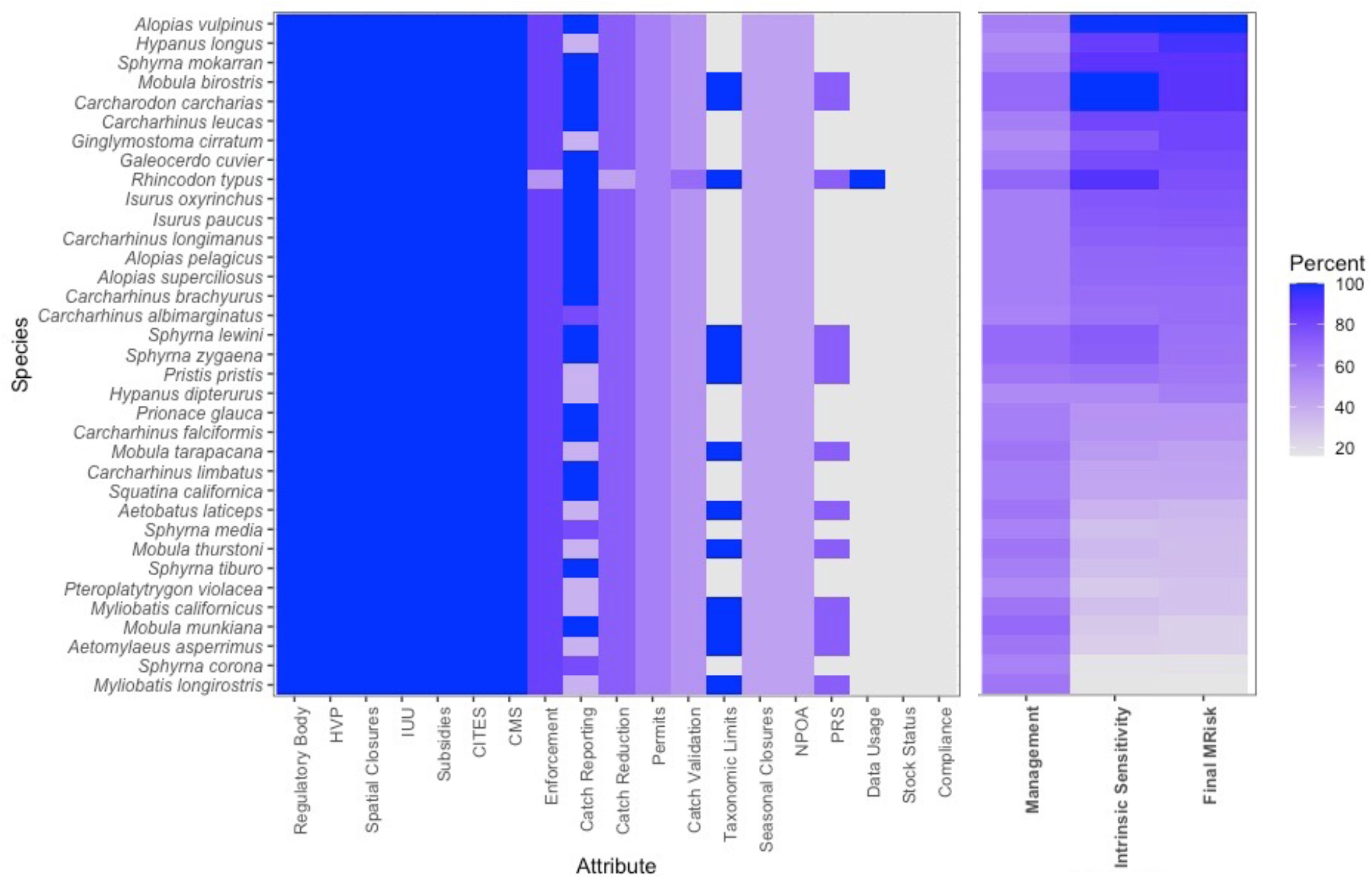
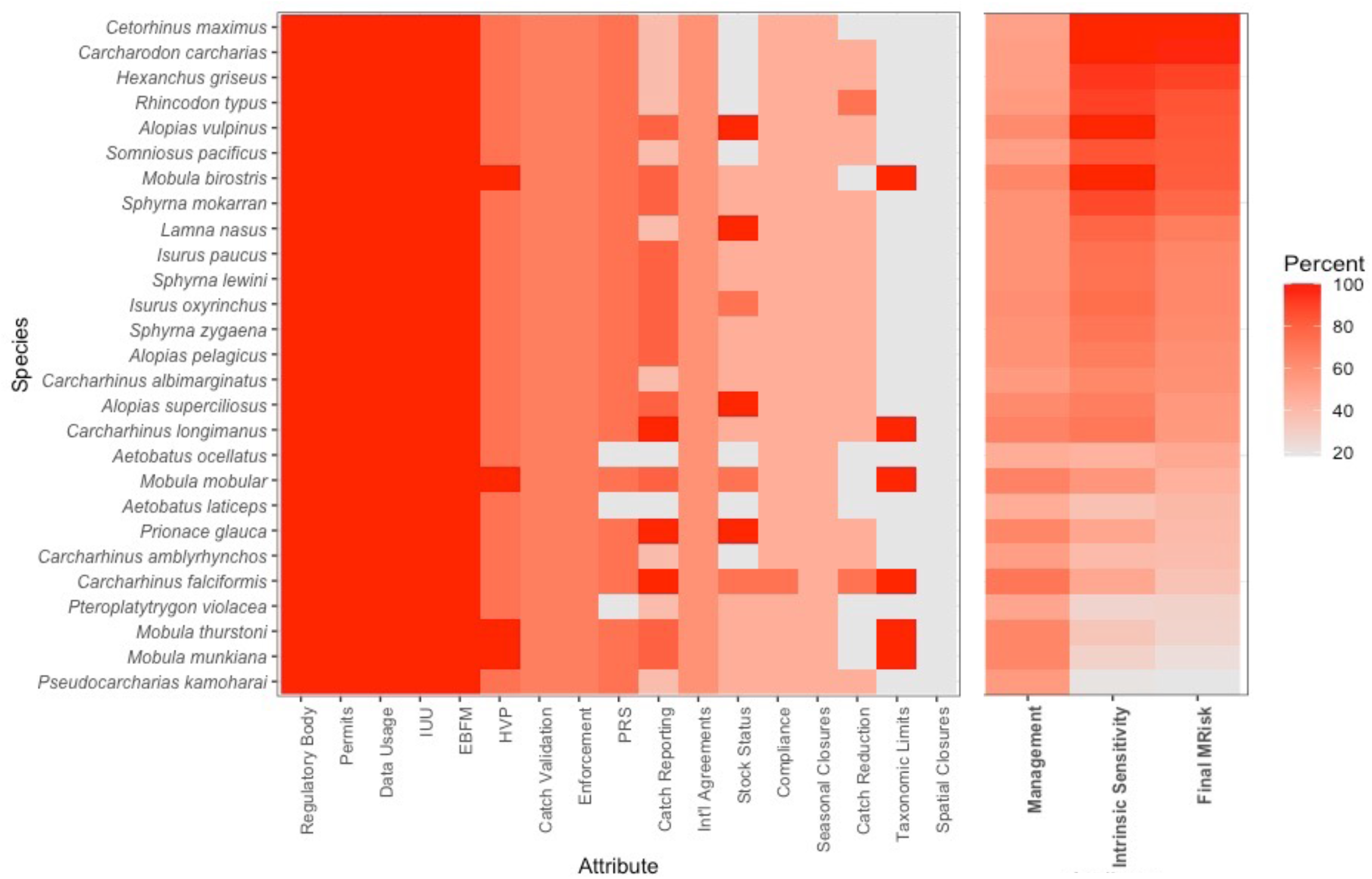
All attribute scores, management scores, intrinsic sensitivity scores, and final M-Risk scores for all species assessed in (a) Ecuador and (b) IATTC. Attribute columns are arranged from highest average score to lowest average score within the management unit. Final M-Risk scores are presented as the percentage of the highest score such that the higher M-Risk scores represent the most at-risk species. No NA values are included in the figure as only relevant attributes for each management unit are included.

##### 3.1.1.3 Management Assessment Discussion

There was little connection between the IUCN status and M-Risk scores with threatened species included in both higher and lower scoring groups. Almost all species with higher scores are listed on both CITES and CMS appendices, indicating clear progress for listed species. All species listed on CITES prior to 2014 were included in the higher scoring group.

Eight CITES and nine CMS-listed species were in the lower scoring group, indicating it takes some time to implement regulations relating to these agreements. This has been seen in other CITES-listed species, like seahorses, which were listed on CITES in 2002, however, Thailand continued exporting high numbers without a positive Non-Detriment Finding until 2016 (Kuo, Laksanawimol, Aylesworth, Foster, & Vincent, 2018). What was more likely to determine a higher score was species charisma. Charismatic species likely to be of high tourism interest are often subject to increased conservation efforts (Albert, Luque, & Courchamp, 2018; Hausmann, Slotow, Fraser, & Di Minin, 2016; McClenachan, Cooper, Carpenter, & Dulvy, 2011), whereas a very large fraction of highly threatened species can go unstudied and unmanaged (Guy et al., 2021). This was apparent in the scores in Ecuador, as all species in the higher scoring group had some type of retention ban or catch limit (**Table S1**). These limits, in addition to CITES and CMS compliance, were likely placed to protect tourism value and public perception.

#### 3.1.2 Case Study 2: IATTC

##### 3.1.2.1 Information Sources

Like assessments for Ecuador, we searched Google and Google Scholar for the same four elements: (i) IATTC fisheries regulations, (ii) specific attribute keywords, (iii) species name (English and Latin binomial), and (iv) species name and specific attribute keywords. For these assessments, we scored both the purse seine and the longline regulations, and the higher score was used. We used one primary and three secondary resources to complete the IATTC management risk assessments. Primarily, we used the IATTC website (www.iattc.org), which provided in-depth documents related to management regulations and fishery knowledge. Additionally, we found information from: (1) the Food and Agriculture Organization of the United Nations website (ww.fao.org), (2) the National Oceanic and Atmospheric Agency website (www.noaa.gov), and (3) the International Scientific Committee for Tuna and Tuna-like Species in the North Pacific Ocean website (isc.fra.go.jp).

##### 3.1.2.2 Management Assessment Results

Scores from the 27 species assessed ranged from 46–71% of an ideal score in the IATTC, with an average score of 58.3% (**Figure 3b; Table S1**). A single species, the Silky Shark (*Carcharhinus falciformis*), stood out with the highest score (71%). Three species, the Pelagic Stingray (*Pteroplatytrygon violacea*) and two eagle ray species (*Aetobatus laticeps and A. ocellatus*) both had scores of 50% or lower (50, 46 and 46%, respectively) and 14 of the 27 species scored between 50-59% (**Figure 3b; Table S1**). Catch reporting was the attribute with the lowest scores for the IATTC because half of the species were only reported to broad taxonomic categories (i.e., “sharks” or “rays”). Only six species assessed had any form of landing limits in place (Silky Shark, Oceanic Whitetip Shark (*Carcharhinus longimanus*), Giant Manta Ray (*Mobula birostris*), and devil rays (*Mobula mobular, M. munkiana*, and *M. thurstoni*), while the rest were only *recommended* to be released once caught. These results show that overall, the IATTC has species-specific regulations in place for only a select few species and the others are subject to marginal management applicable to elasmobranchs. It remains unclear whether these measures can significantly reduce fishing mortality and if the management focus is on the species that are most at-risk to overfishing within the fishery.

##### 3.1.2.3 Management Assessment Discussion

Within the IATTC, there does not appear to be any relationship between global IUCN status and the management assessment scores. Of the 27 species assessed, only five are not currently listed in a threatened category, yet these are not the five with the lowest scores. Similarly, CITES and CMS listings appear to have not had any bearing on the management assessment scores. The Whale Shark has been listed on both CITES (2003) and CMS (App I in 2017, App II in 1999) for much longer than many other sharks and rays and is unquestionably one of the most charismatic species, which we would expect to lead to a higher management score. However, it received a score of only 55% in the IATTC. The score may be lower than expected as there is little interaction between the fishery and whale sharks, thus lesser concern to provide specific legislation for this species. Additionally, where interactions do occur, there is a prohibition on setting nets when a whale shark is sighted (Inter-American Tropical Tuna Commission, 2019). The Silky Shark, which has been more recently listed on both CITES (2017) and CMS (2014) appendices, received the highest score of all species assessed. This surprisingly rapid management response may be due to the large amount of recent attention and market pressure to reduce the plight of Silky Shark caught in fish aggregating devices (FADs) and their susceptibility to purse seines (Duffy et al., 2015; Filmalter, Capello, Deneubourg, Cowley, & Dagorn, 2013; Hutchinson, Itano, Muir, & Holland, 2015). Although the purse-seine tuna fishery catches mostly shark ‘bycatch’, there are several pelagic ray species caught within the fishery. Although CITES listing may not necessarily be considered by RFMOs when discussing regulations, it should be. CITES is a legally binding agreement which covers relevant issues like ‘Introduction From the Sea’, including the high seas that constitute the fishing grounds plied by vessels operating under RFMOs, and countries’ responsibilities for legal chain of custody when CITES-listed species are landed from high seas vessels (Pavitt et al., 2021). With the exception of manta and devil rays, which are no-retention species (Inter-American Tropical Tuna Commission, 2015), there are no ray-specific regulations to reduce their catch. While shark species benefit from more generic legislation including the prohibition of shark lines (which intentionally catch sharks as retained bycatch), a code of practice to increase post-release survival, and fin-to-meat landing ratios to reduce finning, rays are not afforded any additional regulations (Inter-American Tropical Tuna Commission, 2005, 2016). In order to decrease management risk for elasmobranch species in the IATTC, implementing science-based landing limits for the most commonly caught species and including more legislation to evaluate and reduce ray fishing and post-release morality are recommended.

### 3.2 Final M-Risk Scores

The final M-Risk score for a species shows whether the current management is appropriate based on its intrinsic sensitivity (IS). Species with high intrinsic sensitivity will need higher management assessment scores to achieve a similar M-Risk score to lower IS species. As an example, the Great Hammerhead (*Sphyrna mokarran*) has a generation length of 24.8 years for a Relative GL score of 80% and reaches a maximum size of 610 cm total length, which is 95.3% of the maximum sized shark (White Shark – 640 cm), therefore, the combined intrinsic sensitivity score for Great Hammerhead is 87.7% (**Figure 2; Table S1**). Compared to the Pelagic Stingray (*Pteroplatytrygon violacea*), which has a generation length of 6.5 years (*Relative GL of 20%*) and reaches a maximum size of 90 cm disc width, which is 34.6% of the maximum sized classic ray (Spiny Butterfly Ray, *Gymnura altavela* – 260 cm), therefore, the combined intrinsic sensitivity score for Pelagic Stingray is 27.3%. Although both the Great Hammerhead and Pelagic Stingray received similar scores in their management assessments for both Ecuador and the IATTC, when combined with their intrinsic sensitivity scores, the final Vulnerability is vastly different. The Great Hammerhead is vastly undermanaged for its life history compared to the Pelagic Stingray (**Figure 3; Table S1**). Although the Pelagic Stingray scored lower in the management assessment for the IATTC than the Great Hammerhead (50.0% and 58.9%, respectively), the final M-Risk scores show it is almost one-third at risk due to undermanagement in the IATTC than the Great Hammerhead due to its lower intrinsic sensitivity (M-Risk scores of 0.51 and 1.50, respectively). In Ecuador, the Pelagic Stingray is similarly approximately one-third as at-risk compared to Great Hammerhead (0.55 and 1.49, respectively). These scores describe the true risk to undermanagement by each of these species based on our current understanding of these species and their populations.

## 4. DISCUSSION

Here, we presented a framework for a management risk assessment (M-Risk) that is designed: (1) for assessing different management approaches for individual species *within* a management unit (of country or RFMO), (2) for comparing the efficacy of shark and ray management *across* different management units globally, and (3) for comparing management efficacy of a single species in all management units it is found. We also summarise the two main findings from this proof-of-concept. Firstly, different shark and ray species within a single management unit have management assessment scores that range from ∼50-70% of a score that would be consistent with the ‘ideal’ management in the two fisheries assessed (Ecuador and the IATTC). Second, when accounting for species’ intrinsic sensitivities, the final M-Risk scores (Vulnerability in Equation 5) best show which species are most at risk due to management deficiencies.

Intrinsic sensitivity of each species is important and particularly beneficial to our understanding of risk differences for species caught in poorly managed fisheries that have low intrinsic sensitivity, compared to high-risk species in adequately managed fisheries. Incorporating species-specific sensitivities to this analysis shows the nuance of managing sharks and rays, which are frequently viewed as a large group with similar attributes. With this approach, the difficulty of having good management for higher risk species is demonstrated as those with high intrinsic sensitivity scores will always have a higher final M-Risk score than species with lower intrinsic sensitivities. Upon completion of a wider array of countries with varying management regimes, we will be able to better assign final

M-Risk scores to either low, medium, or high-risk groupings. This will provide an understanding and priority setting for which species, in which management units, are most at risk of overexploitation and at what M-Risk score intervention should begin. Additionally, after more management units are scored, we will evaluate the strength of pairwise correlations between attribute scores and/or country and species traits. This will allow us to ask questions such as, “do countries with higher Human Development Indices always score higher in the ‘Country Attributes’?” and “in fisheries with a high score in the Catch Validation attribute, do they always receive a high score for Enforcement Methods?” The answers to these questions may uncover causal relationships that provide clear paths to improving fisheries management. Next, we explore the differences in scores across management units, how our M-Risk approach compares to other PSAs, and future directions for this work.

Two management units (IATTC and Ecuador) were assessed using the M-Risk framework. Once additional management units are assessed, countries will be compared to other countries and RFMOs can all be compared to one another. In describing the applications of this risk assessment framework, this discussion will assume these two management units are directly comparable. Despite both receiving similar scores for species in their respective jurisdictions, the scores for specific attribute categories differed. For example, the IATTC species, on average, scored 30% higher in the “Management System” attributes than species in Ecuador (73% and 44%, respectively). However, in the “Fishing Practices and Catch” attributes, species in Ecuador scored, on average, 20% higher than those in the IATTC (59% and 37%, respectively; **Figure 3; Table S1**). Based on these results, each management unit could benefit from studying regulations in the other to improve their own shark and ray management. Comparing management units using the same framework not only enables a level playing field, but also allows for management units with lower scores to more easily identify areas to improve their management efficacy. Management units lacking sufficient funding to complete their own fisheries research into which regulations have the greatest impact on shark and ray fishing mortality can learn from or borrow legislation from those with highly effective management strategies. This can also apply to sections of management that are lacking in sufficient shark and ray management. If adopted, this would increase the efficacy of global fisheries management for sharks and rays overall. A limitation of this type of assessment is it does not consider compliance with legislation. This includes compliance by the fishers, fisheries officers, and managers who are policing the regulations and assumes countries are meeting their obligations to agreements to which they are signatories, including CITES and for RFMOs. Without effective compliance and enforcement of the fisheries regulations, the management scores we have assigned for each fishery are a best-case scenario and likely over-estimate the management effectiveness. Since these assessments are intended to be completed by external assessors using publicly available data, the scores should be considered minimum possible scores for each management unit if all regulations are not readily available online and consequently, our approach is precautionary.

Future directions for the M-Risk framework may include a capacity to weight attributes and include an evaluation of the exposure component. For this global comparison, weighting attributes is not appropriate as the goal is to compare across all fishery types, gears, countries, etc. For example, in Bangladesh the commercial and artisanal sectors operate with different goals and budgets and in different areas (Islam et al., 2017; Kumar et al., 2019). Therefore, an attribute that may be weighted higher for assessing the commercial fisheries may not make sense to be weighted that way when assessing subsistence fisheries. There are methods available to assign post-hoc weightings to attributes that remove the subjectivity of determining “importance” (Chen, 2019). Post-hoc weighting of attributes may be applied when using the management assessment framework to suit a particular goal. However, as the framework is designed for ongoing use, post-hoc weightings may continue to change as more assessments are completed. In our current assessments, exposure has been treated as a binary variable, despite normally ranging from 0–1, because we are assessing species in trade, therefore, they are assumed to be highly exposed. However, if information about the fishing grounds and depths of species becomes easier to acquire, there is potential to include exposure in future M-Risk work. Additionally, this framework does not consider cumulative impacts due to multiple fisheries or threats, including climate change, environmental modifications and other anthropogenic hazards, that may significantly increase overall risk to some species (Walker et al., 2021). However, this framework could be readily adapted to incorporate vulnerability to climate change through the addition of Equation 3, section 2.2.12 of Walker et al. (2021). We have not explored the implications of multiple assessments through time, however, anticipate this could provide valuable indications of management improvement or decline, given the assessments are completed against the same criteria in the same manner.

The results of this proof of concept provide a basis for further investigation of shark and ray management globally. With a larger number of management units assessed, we will have a better understanding of the efficacy of shark and ray management globally and which management units or species are most at risk due to undermanagement based on their intrinsic sensitivity. The M-Risk assessment tool may be a step to identify species-at-risk and appropriate management to implement that would lower risk to those species. Currently, international agreements like CITES and CMS are in place once species have been depleted. However, identifying species-at-risk to overexploitation and implementing management to rebuild and achieve sustainable fisheries and avoid population collapses, may reduce the need for some species to be included in these international agreements in the first place as maintenance is simpler than recovery.

## ACKNOWLEDGEMENTS

The authors thank Heather Patterson for her input of attributes and their scoring. We thank Thomasina Oldfield and Dulvy lab members for comments on manuscript drafts. Additionally, we would like to thank Ian Gregor for help with coding the graphs. We are grateful to the three reviewers and editor who provided valuable comments that improved this manuscript. This work was supported by a grant to GS from the Shark Conservation Fund, a philanthropic collaborative that pools expertise and resources to meet the threats facing the world’s sharks and rays. The Shark Conservation Fund is a project of Rockefeller Philanthropy Advisors. NKD was supported by the Discovery and Accelerator grants from Natural Science and Engineering Research Council and the Canada Research Chair program. The authors have no conflicts of interest to declare.

## DATA AVAILABILITY STATEMENT

Individual attribute scores for each species in each management unit, and the management documents used for scoring are available in the Supplementary Information (**Data S2 and Data S3**, respectively).

## Notes

### Competing Interest Statement

The authors have declared no competing interest.

