## Supplementary Information, Table S1 for "M-Risk: A framework for assessing global fisheries management efficacy of sharks, rays, and chimaeras"

**Spreadsheets:**

**Data S1.** Full list of species included in the larger M-Risk project, including whether they are considered a group or not, relative maximum size, relative generation length, and overall relative intrinsic sensitivity.

**Data S2.** Individual scores for each attribute of each species in each management unit.

**Data S3.** List of resources used to score each attribute for both the IATTC and Ecuador.

**Supplementary Information 1**

**Attribute Value Statements**

**MANAGEMENT SYSTEM**

***1.***	***Regulatory Body in Place***

Having a regulatory body allows fisheries to be managed. The regulatory body includes the views of fisheries managers, environmental groups, fishermen, and scientists, among others. This ensures that all invested parties have a say in the regulations made in the fishery. Without a regulatory body, a fishery is unable to adequately monitor catch. If this body does not meet regularly, changes in stock status or other stuff may not be addressed in a timely fashion. The score for this attribute will remain the same for all species assessed within a management unit.

0.	No regulatory body in place

1.	Regulatory body that meets less than once every two years

2.	Regulatory body that meets at least once every two years

3.	Regulatory body that meets at least once every year

***2.***	***Permits***

Having permits within a fishery enables managers to determine the number of fishers and allows for easy distribution of new legislation to those that are required to abide by it. Once permits are required, actual fishing pressure can be more accurately calculated. Having a permit that is associated with some form of fishing limit (ITQs or TAC) will enable fisheries managers to better control fishing mortality and adjust if overfishing is occurring. The score for this attribute will remain the same for all species assessed within a management unit.

0.	No permits required

1.	Permits / vessel registry required

2.	Permits with ITQ / TAC / trawl hours / some form of limit associated with the permit

***3.***	***Stock Status / Risk Assessments***

Collecting information regarding stock status is important to ensure overfishing is not occurring and that a population is not overfished. Once information is collected, stock status can be used to assess risk to overfishing. These assessments provide feedback to managers on the sustainability of the fishery and the ecosystem in which the fishery is operating. These assessments can expose areas of the fishery that require further research or can show that there is no need to complete more expensive assessments. In many cases, collating this information and performing an in-depth risk assessment is costly and time consuming, thus it is impractical. If basic risk assessments are performed and indicate there is no need for further assessments, stock status can be assumed to be healthy. In fisheries with the capacity to perform more in-depth stock assessments, results can be useful for smaller, adjacent fisheries. The score for this attribute may differ for each species in the management unit.

0.	No stock status / risk assessments performed

1.	Basic risk assessments performed showing high risk with no follow up assessments performed OR no provisions made for high risk species of elasmobranchs

2.	In depth assessments performed show overfishing is occurring OR basic assessment performed and some provisions made for high risk elasmobranchs

3.	In depth stock assessments show that overfishing is not occurring OR basic risk assessments performed showing no need for further assessments on the species (low risk fishery)

***4.***	***Sustainable Fishing Determination / Data Usage***

The method used for a fishery to determine the total allowable effort or catch (TAE / TAC) of their target species can inform what capacity the fishery has to manage its bycatch species. When determining TACC or total allowable effort, the method used will inform how accurate and safe the allowances are. With more data collected and available, more precise analyses can be done and, therefore, there is a better idea of a sustainable level of fishing mortality. This is measured for the target species that receives the lowest score, whether or not that is an elasmobranch. The method for determining sustainable fishing limits is telling of the overall management and resources available within the management unit. The score for this attribute will remain the same for all species assessed within a management unit.

0.	None

1.	Based on ERA results, estimations, or previous TACC/ effort

2.	Based on CPUE

3.	Based on stock assessments

***5.***	***Efforts to Reduce Catch***

In fisheries in which the species being assessed is not targeted, efforts to reduce bycatch should be in place to lower overall fishing mortality of the species. Methods to reduce catch of sharks and rays include bycatch reduction devices (BRDs) like turtle excluder devices, depth restrictions, trigger points, move-on limits. BRDs are frequently used in trawl fisheries and can significantly reduce the catch of larger elasmobranchs (Griffiths, Brewer, Heales, Milton, & Stobutzki, 2006). Depth restrictions have also been shown to reduce elasmobranch bycatch and in some instances, increase commercial value (Clarke et al., 2015). Trigger points allow fisheries managers to identify a sustainable maximum catch and then stop commercial fishing operations once this has been hit for the year. Move-on limits are used in some fisheries and can be effective for species that aggregate. These limits require vessels to move a predetermined distance from the last point their gear was deployed if a certain level of bycatch is met to ensure another deployment does not completely remove the species in question from the area (Federal Fisheries Council, 2013; Nowara et al., 2017). The score for this attribute may differ for each species in the management unit.

0.	No efforts in place to reduce catch

1.	One catch reduction method in place

2.	Two catch reduction methods in place OR one scientifically based catch reduction method

3.	Three or more catch reduction methods in place OR two or more scientifically based catch reduction methods

**FISHING PRACTICES / CATCH**

***6.***	***Taxonomic Resolution of Landing Limits***

Having science-based landing limits in place is the only direct method to impact fishing mortality of a species. Landing limits can be set based on previous catch, ERAs, scientific advice, or stock assessments, among other methods. Having landing limits for all sharks or rays is a first step in limiting the amount that can be taken. However, this limit may not be beneficial to some threatened species. For example, catching 10 tonnes of the Endangered Whale Shark (*Rhincodon typus*) would have a much bigger impact on the ecosystem than catching 10 tonnes of a Least Concern species, like the Milk Shark (*Rhizoprionodon acutus*). Species-specific catch limits are required to maintain a healthy elasmobranch community. The score for this attribute may differ for each species in the management unit.

0. No Landing limits set

1. Landing limit for CLASS / SUBCLASS “sharks / rays”

2. Landing limit for ORDER / FAMILY “hammerheads, thresher, deepwater sharks, etc.”

3. Species-specific landing limits OR no retention including the species being assessed

**7.**	***Removal of High Value Products (HVP)***

Many elasmobranchs have high value products (HVPs) that can be easily removed at sea. Vessel space is limited, thus fishers may remove HVPs at sea and discard the rest of the carcass in order to save valuable space and maximise their profitability. HVPs include fins, livers, *Mobula* gill plates, and rostra, among others. HVP at-sea removals have the potential to curb our ability to monitor species-specific catches. Fisheries have a few options to reduce HVP removal. These regulations ensure that entire carcasses are not wasted and when enforced, likely reduce cryptic mortality. The score for this attribute may differ for each species in the management unit.

0.	No regulations against removing any high value products at sea

1.	Fin to carcass ratios on landed sharks OR only regulations relating to removal of some HVPs relevant to the management unit

2.	Fin to meat ratios on landed sharks OR regulations relating to removal of all HVPs relevant to the management unit

3.	Fins naturally attached on landed sharks AND regulations relating to at-sea removal of all HVPs relevant to the management unit

***8.***	***Seasonal Closures***

Seasonal closures in fishery grounds allow for populations to naturally grow in the absence of fishing pressure. These closures are often implemented to allow for target species to spawn, breed, and/or pup. The reduction of fishing pressure can either incidentally or directly reduce fishing mortality of elasmobranchs, depending on the intention. In some areas, seasonal closures are in place, however, due to weather there would be no fishing operations at these times so they do not reduce fishing mortality further. The score for this attribute may differ for each species in the management unit.

0. No closures OR seasonal closure unlikely to curb fishing mortality (i.e. closed for monsoon season)

1. Seasonal closure for target species (non-elasmobranch) spawning / protection OR overall reduction in catch

2. Seasonal closure in place with the intention of reducing overall elasmobranch mortality

3. Seasonal closure for the benefit of the species being assessed (i.e. breeding/pupping grounds during season)

***9.***	***Spatial Closures***

Spatial closures can be put in place to protect important habitats for certain species, or a portion of a species’ range. These closures are often implemented in known sensitive locations for target species including areas for spawning, breeding, and/or pupping. The reduction of fishing pressure can either incidentally or directly reduce fishing mortality of elasmobranchs, depending on the intention. In some cases, spatial closures are put in place to benefit bycatch species, including some elasmobranchs. The score for this attribute may differ for each species in the management unit.

0.	No spatial closures

1.	Spatial closures in place for non-elasmobranchs

2.	Spatial closures in place for elasmobranchs but not the species being assessed

3.	Spatial closures in place for the species being assessed

***10.*** ***Post-Release Survival***

Post-release survival (PRS) estimates the likelihood of survival of animals caught by fishing gears and released back to the ocean. Estimates are based on condition of individuals as they are released. These estimates can be substantiated with tagging studies. Estimates are important because, when known, there can be measures put in place to increase the survival. Additionally, if there is high PRS, sustainability is less of a concern. As well as an estimate, measures can be taken to increase the likelihood of bycatch survival. These include circle hooks, which are more likely to hook a shark in the mouth than a deep hooking, and hoppers that reduce the time spent out of water while being sorted on the boat. Additionally, some fisheries include a code of practice on how to handle sharks in a manner that increases their survival once released. The score for this attribute may differ for each species in the management unit.

0.	No indication of PRS estimate or measures to increase PRS

1.	A post-release survival estimate

2.	Measures to increase post-release survival (through fishing gears modification i.e. circle hooks, hoppers, and or modification of handling practice e.g. Code of practice on shark handling to increase post-release survival)

3.	Both a PRS estimate and measures to increase PRS

**COMPLIANCE, MONITORING, AND ENFORCEMENT**

***11. Catch Reporting***

Collecting catch information including ALL non-retained catch is important in understanding potential fishing mortality of a species. Species-specific information is best to fully understand the impact that a fishery has on a given species. As some species are more susceptible to post-release mortality, their catch levels may be unsustainable even without retention of the animals. For countries, when data cannot be found for the specific fishery, this will be scored by the most recent taxonomic level of reporting to FAO. Scores are only given if there are numbers associated with the species catch (i.e. a list of species caught in a fishery without the amount of each species caught is not considered). The score for this attribute may differ for each species in the management unit.

0.	No reporting of elasmobranch catch

1.	Catch reported to broad categories (i.e. “sharks” “elasmobranchs” or “rays”) OR list of species listed with no associated numbers of actual catch

2.	Catch reporting to narrow categories (i.e. “mackerel sharks,” “reef sharks,” “deepwater sharks,” “whaler sharks,” etc.)

3.	Catch reporting of similar/related species grouped (“deepwater dogfishes,” “gulper sharks,” “mako,” “hammerhead,” etc.)

4.	Species-specific catch reporting

***12.*** ***Illegal, Unregulated, and Unreported (IUU) Fishing***

Illegal, unregulated and unreported (IUU) fishing is hazardous as the complete fishing mortality of each species is not known. While IUU fishing cannot be stopped 100% due to the vast space of the ocean and relatively small size of the boats, it can be considered in management decisions including quota considerations and increased patrolling in areas with frequent IUU. The score for this attribute will remain the same for all species assessed within a management unit.

0.	IUU fishing a problem but not recognised in management documents

1.	IUU fishing a problem and acknowledged in management documents

2.	IUU fishing a problem and management arrangements reflect the impact IUU has in the management unit

3.	IUU fishing not a problem OR management arrangements to address IUU appear successful OR a signatory to the Agreement on Port State Measures (PSMA)

***13.*** ***Compliance Regime***

Fisheries management includes both generic and species-specific measures in order to control for fishing mortality. Some generic measures include limited entry (licensing system), gear restrictions, permanent area closures, etc. These regulations often will positively affect both target and non-target species populations by reducing fishing pressure and providing refuge areas. Species-specific measures include size limits, gender restrictions, move-on provisions, etc. These are commonly put in place for frequent bycatch species due to the impact fishing pressure may have on their populations. Not all of these regulations are relevant to the fishing mortality of each species assessed, therefore, they will be assessed based on whether the generic and/or species-specific measures are likely to reduce fishing mortality. The score for this attribute may differ for each species in the management unit.

0.	No species-specific measures in place / no relevant compliance measures in place

1.	Compliance measures in place unlikely to significantly reduce fishing mortality

2.	Compliance measures in place likely to reduce fishing mortality

3.	Species-specific measures in place that have proven to reduce fishing mortality

***14.*** ***Catch Validation***

All fisheries need some form of monitoring of fishing activities. Monitoring allows for managers to determine the level of activity and compliance within the fishery. Forms of monitoring include logbooks, various electronic monitoring systems, and observer programs or video observer programs (only when recordings of ALL catch of elasmobranchs are available). The score for this attribute will remain the same for all species assessed within a management unit.

* Observer program must be permanently in place to receive the points. Some fisheries put observers in place sporadically, therefore, only those fisheries that have observers every year will be considered. Fisheries with observer programs that do not operate each year will not receive any points for the program.

One point each for:

·	Self-reporting through logbooks

·	EMS (if there is video that covers 100% of the hauling of equipment)

·	Observer program* with less than 33% coverage (1 point)

·	Observer program* with up to 67% coverage (2 points)

·	Observer program* with 100% coverage (3 points)

***15.*** ***Enforcement Methods***

Methods of enforcement are measures taken that deter fishermen from breaking regulations. Enforcement penalties should provide higher incentive for fishermen to abide by regulations than they would have to fish illegally. Enforcement can include fines, criminal records, even jail-time, depending on the severity of the infraction. Fisheries have several ways of acquiring information that will lead to punishments. More ways to find those breaking rules will lead to more enforcement and better deter those performing illegal activities, therefore, this attribute is measured in the methods taken of finding those illegally fishing. The score for this attribute will remain the same for all species assessed within a management unit.

One point each for:

·	Use of electronic surveillance (e.g. drones, and/or planes, and/or VMS) to fine boats that are outside of regulations

·	Boarding of vessels to ensure compliance

·	Port checks for under/ oversized catch and/or illegal species

·	Logbook validation

·	Monitoring and/or weighing of catch unloading

**COUNTRY INDICATORS**

These are to be used when assessing fisheries from a single country (no RFMOs or RFBs)

***16.*** ***Subsidies***

Subsidies are provided to fishermen to ensure they are profitable. Some subsidies ensure profitability by enhancing the capacity of fishermen to increase their catch, often at the expense of sustainability. These deleterious “capacity-enhancing” subsidies include fishery development, boat, market infrastructure, port, tax exemptions, access, and fuel subsidies. On the contrary, some subsidies enable fishermen to continue fishing sustainably without a loss of profit. Beneficial subsidies include those for fisheries management, research and development, and marine protected area implementation and management. The score for this attribute will remain the same for all species assessed within a management unit.

0.	Amount given for deleterious subsidies more than double the amount for beneficial subsidies

1.	Amount given for deleterious subsidies higher than for beneficial subsidies

2.	Amount given for beneficial subsidies higher than for deleterious subsidies

3.	Amount given for beneficial subsidies more than double the amount given for deleterious subsidies

***17.*** ***NPOA-Sharks and its Effectiveness***

Having an NPOA for sharks and rays shows a country is interested in the welfare of its shark populations. However, if certain NPOA objectives are not addressed, then it is unlikely to provide effective solutions for mitigating fishing mortality of sharks. Ten objectives were proposed by Davis and Worm (2013) as the minimum content to include. The score for this attribute will remain the same for all species assessed within a management unit.

0. 0-3 objectives addressed in NPOA

1. 4-6 objectives addressed in NPOA

2. 7-9 objectives addressed in NPOA

3. All 10 objectives addressed in NPOA

***18.*** ***CITES Member***

Is the country a member country of CITES? If so, do they have any reservations on listed chondrichthyan species? Being a member country of CITES indicates an overall interest in conservation of at-risk species. However, if there are reservations in place for species that should have additional management, it negates the impact of being a CITES member. The score for this attribute may differ for each species in the management unit.

0.	Not a member CITES

1.	CITES member with reservations on >50% of listed chondrichthyan species OR a reservation on the species being assessed

2.	CITES member with reservations on <50% of listed chondrichthyan species

3.	CITES member with no reservations on chondrichthyan species

***19.*** ***CMS Member***

Is the country a member country of CMS or a signatory on the MOU-Sharks? The score for this attribute will remain the same for all species assessed within a management unit.

0.	Not a member of CMS

1.	CMS member OR MOU-Sharks Signatory

2.	CMS member AND MOU-Sharks Signatory

**RFMO INDICATORS**

These are to be used when assessing RFMOs or RFBs.

**20.** **All Countries Involved are CITES / CMS Member**

Are member countries or non-member contracted parties that fish within the RFMO members of CITES / CMS? If a CITES member, do they have any reservations on the species being assessed? The score for this attribute may differ for each species in the management unit.

0.	No member countries are part of CITES or CMS

1.	<50% of member countries part of CITES and/or CMS

2.	>=50% of member countries part of CITES and/or CMS

3.	100% of member countries part of at least one of CITES with no reservations on the species being assessed or 100% members of CMS

4.	100% of member countries part of both CITES with no reservations on the species being assessed and of CMS

**21.** **Ecosystem Based Fisheries Management Implementation**

How well progressed is the RFMO with implementation of ecosystem-based fisheries management? Scored using results from Juan-Jordá et al. 2017 where a score of slight or no progress = 0, moderate progress by scientific committee = 1, full progress by only the scientific committee = 2, slight progress by commission = 3, moderate progress by commission = 4, full progress by commission = 5. All 20 ecological component scores will be averaged for the RFMO. The score for this attribute will remain the same for all species assessed within a management unit.

0.	Average score of <2

1.	Average score 2.1-3

2.	Average score 3.1-4

3.	Average score 4.1 +

**Supplementary Information 2**

**Intrinsic Sensitivity Measures**

Intrinsic sensitivity can be measured through a combination of many variables including age-at-maturity, length-at-maturity, longevity, maximum size, fecundity, reproductive strategy, and trophic level (Musick, 1999; Oldfield, Outhwaite, Goodman, & Sant, 2012). Preferentially, we would have used r_max_ as a measure of Intrinsic Sensitivity (Pardo, Cooper, Reynolds, & Dulvy, 2018). However, these data are not available for most shark and ray species. Without r_max_ data, generation length (GL) would be the best indicator, if these values were available for each species. Where GL data was not available, we used the values estimated in the species’ IUCN Red List Assessment or estimated in a similar manner, through congener species. We then assigned a confidence to the GLs for each species (low, medium, or high). For species without species-specific data, where GL was estimated and scaled based on a congener species, confidence was scored as low (65.3%; n = 213 of 326). Some species had estimated species-specific data used to determine GLs, these were also assigned low confidence (1.6%; n = 5), for a total of 66.9% of species having low confidence GLs. As we aggregated generation lengths to 5-year bins due to uncertainty, we assigned species with GL estimates that spanned multiple bins medium confidence as we precautionarily used the highest GL value (14.1%; n = 46). We assigned high confidence to species with species-specific data that led to a GL estimate within a single 5-year bin (19.0%; n = 62).

The low confidence in generation length values can best be illustrated when looking through the Etmopterus genus. There are species-specific data for *Etmopterus spinax* (7.8 year GL)*, E. pusillus* (12.5 yrs)*, E. princeps* (21 yrs)*,* and *E. granulosus* (43.5 yrs), which are all deepwater species in the same genus encompassing at least 38 species. The latter two species were only assigned medium confidence as their GLs were estimated based on suspected age-at-maturity and/or longevity, not validated age data (Irvine, Stevens, & Laurenson, 2006; Kulka et al., 2020). The rest of the Etmopterus species considered (n = 34) have GLs estimated based on *E. pusillus* (the longer GL of the two species with high confidence), however, with extreme uncertainty.

Intrinsic sensitivity is further complicated when comparing species with different depth ranges. Deepwater species tend to have lower productivity, and therefore, higher intrinsic sensitivities (Rigby & Simpfendorfer, 2015). Shark productivity (i.e. intrinsic rate of population increase *r_max_*) decreases with increasing depth, despite no corresponding increase in size (Georgeson et al., 2020). However, in skates, there is an increase in size with depth, therefore, the corresponding relationship between productivity and depth cannot be disentangled from size (Georgeson et al., 2020). Although the slope of the relationship may be different, there is a significant positive relationship between female size-at-maturity and maximum body length in deepwater shark and ray species (Rigby & Simpfendorfer, 2015), and therefore generation length should increase with body length. This bolsters our use of maximum body length in addition to generation length when calculating Intrinsic Sensitivity. However, we do note that there may be a relationship between the two variables.

**Generation Length and Maximum Body Size Relationship**

The relationship between generation length and relative maximum body size was tested using a simple linear regression model using R package lme4 (Bates et al., 2019). Generation length (GL) was included in the model as number of years, rather than the 5-year binned values used for estimating Intrinsic Sensitivity. Maximum body length was included as a relative parameter, scaled based on the largest species, as per the Intrinsic Sensitivity measurement. Rosner’s test for outliers was performed using R package EnvStats (Millard & Kowarik, 2022) and subsequently two species were excluded Greenland Shark *Somniosus microcephalus* and Pacific Sleeper Shark *Somniosus pacificus* (R_i_ = 11.64 and 6.74, λ_i_ = 3.75), both having generation lengths >75 years. Linear models where relative maximum body length was tested against the interaction of GL and GL confidence (as low, medium, or high) showed a significant relationship between maximum body length and GLs with low and medium confidence (*t*-value = 6.27 and 2.35, degrees of freedom *d.f.* = 216 and 44, *p* = <0.001 and 0.023, respectively). However, no significant relationship between maximum body length and GL was found when using the high confidence GLs (*t*-value = 1.60, df = 60, *p* = 0.114; **SI2 – Figure 1**). Species with low and medium confidence GLs had significantly different slopes from each other as per a least-squares pairwise comparison (R package – lsmeans; (Lenth, 2016)) (*t*.ratio = 2.38, *d.f.* = 318, *p* = 0.047). The high confidence GL slope did not significantly differ from either the low or medium confidence GL slopes (*t*.ratio = 1.88 and -0.29, *d.f*. = 318, *p* = 0.146 and 0.955, respectively)(**SI2 – Figure 1**). Species with low confidence GLs had significantly different intercept values to species with medium and high confidence GLS (*t*.ratio = -3.67 and -3.07, *d.f.* = 318, *p* = 0.001 and 0.007, respectively). Species with medium and high confidence GLs did not significantly differ in intercept values (*t.*ratio = 0.72, *d.f.* = 318, *p* = 0.752). These results were expected as high confidence GL species were those in which GL was calculated based on species-specific age-at-maturity and longevity, whereas species with lower confidence GLs were those with GLs that were estimated and scaled to size, based on data from congeners. By inherently scaling the GL with size, a significant relationship was expected for these species. These results bolster our decision to calculate intrinsic sensitivity using both GL and maximum body length, rather than a single metric.

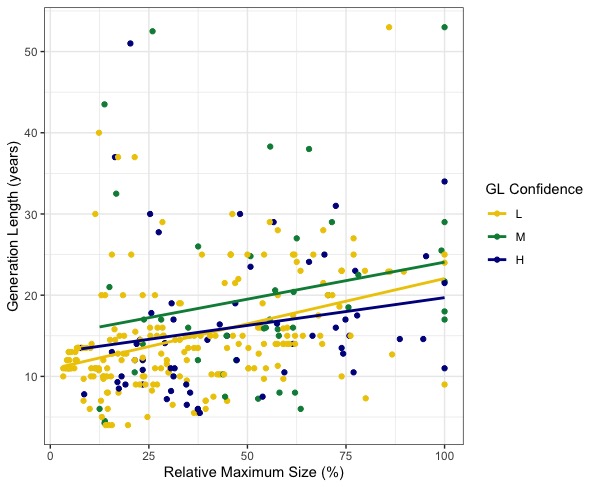

**SI2 - Figure 1.** Relationship between relative maximum size (as a percentage of the largest species) and generation length in years. There was a significant correlation between the two variables for generation length values with low and medium confidence, but not those with high confidence. The slope for species with low confidence was significantly steeper than for species with medium confidence. All other pairwise slope comparisons were not significantly different.
